# Formulation of Dry Powders of Vaccines Containing MF59 or AddaVax by Thin-Film Freeze-Drying

**DOI:** 10.1101/2021.10.21.465307

**Authors:** Khaled AboulFotouh, Naoko Uno, Haiyue Xu, Chaeho Moon, Sawittree Sahakijpijarn, Dale J. Christensen, Gregory J. Davenport, Chris Cano, Ted M Ross, Robert O. Williams, Zhengrong Cui

## Abstract

Oil-in-water (O/W) nanoemulsion-based vaccine adjuvants such as MF59^®^ are often used in seasonal and pandemic influenza vaccines. However, vaccines containing nanoemulsions require cold chain for storage and are sensitive to accidental freezing. We explored the feasibility of developing dry powders of vaccines adjuvanted with MF59 or AddaVax™, a preclinical grade nanoemulsion that has the same composition and droplet size as MF59, by thin-film freeze-drying (TFFD). AddaVax alone was successfully converted from a liquid to dry powders by TFFD using trehalose as a stabilizing agent while maintaining the droplet size distribution of the AddaVax when reconstituted, whereas subjecting the same AddaVax composition to conventional shelf freeze-drying led to significant aggregation or fusion. TFFD was then applied to convert liquid AddaVax-adjuvanted vaccines containing either model antigens such as ovalbumin and lysozyme, mono-, bi-, and tri-valent recombinant hemagglutinin (rHA) protein-based H1 and/or H3 (universal) influenza vaccine candidates, as well as the MF59-containing Fluad^®^ Quadrivalent influenza vaccine to dry powders. Antigens, stabilizing agents, and buffer showed different effects on the physical properties of the vaccines (*e*.*g*., mean particle size and particle size distribution) after subjected to TFFD, but the integrity and hemagglutination activity of the rHA antigens did not significantly change and the immunogenicity of reconstituted influenza vaccine candidates was preserved when evaluated in BALB/c mice. The vaccine dry powder was not sensititve to repeated freezing-and-thawing, in contrast to its liquid counterpart. It is concluded that TFFD can be applied to convert vaccines containing MF59 or an nanoemulsion with the same composition and droplet size as MF59 from liquid to dry powders while maintaining the immunogencity of the vaccines, and it may be used to prepare dry powders of multivalent universal influenza vaccines.

## 1. Introduction

O/W nanoemulsion-based vaccine adjuvants such as MF59 and AS03 can increase and broaden the immune responses induced by vaccines and spare vaccine doses [1-3]. MF59 in a Novartis’ proprietary vaccine adjuvant. It is a squalene oil (4.3% *w/v*) in citrate buffer nanoemulsion stabilized with Tween 80 (0.5% *w/v*) and Span 85 (0.5% *w/v*)), with a mean droplet size of 160 nm [4]. MF59 act by creating a transient immunocompetent environment locally at the site of injection with subsequent recruitment of key immune cells that transport the antigen and adjuvant to local lymph nodes where immune responses are induced [5]. The adjuvant effect of MF59 is maintained in conditions associated with CD4^+^ T cell deficiency, which may explain its effectiveness in broad population and immunocompromised patients [4]. The immunostimulatory activity of MF59 is a property of the nanoemulsion; only the fully formulated MF59 nanoemulsion, not the individual components, can help activate immune responses [6]. Thus, the appropriate formulation composition and physical properties are crucial for MF59 to exert an adjuvant effect. AS03 has a composition and a mean droplet size like that of MF59; however, AS03 contains α-tocopherol, in addition to squalene [7-9]. The presence of α-tocopherol in AS03 is necessary for it to help induce high antibody titers [8].

MF59 is commonly used in seasonal and pandemic influenza vaccines [4]. It enhances the efficacy of these vaccines by eliciting hemagglutination inhibition antibodies as well as memory T and B cells against influenza viruses including antigenic “drift” and “shift” [10]. Examples of MF59-adjuvanted influenza vaccines include the Audenz™(an influenza A (H5N1) monovalent vaccine) and the Fluad^®^ Quadrivalent (an influenza vaccine against influenza virus subtypes A and B). MF59 was also used in pandemic H1N1 vaccines (*e*.*g*., Arepanrix™, Celtura^®^ and Focetria^®^) licensed in many countries during the 2009 H1N1 pandemic [4]. Addtionally, MF59-adjuvanted vaccines against various other infections are currently in clinical trials (*clinicaltrials*.*gov*). AS03 is used in the FDA-approved influenza A (H5N1) monovalent vaccine manufactured by ID Biomedical Corporation of Quebec (QC, Canada). AS03 was also used in Pandemrix™ and Arepanrix™ that were approved during the influenza A (H1N1) pandmic in 2009, though the vaccines were removed from the markets after the pandemic [11].

Nanoemulsion-adjuvanted vaccines are marketed as injectables, either in pre-filled syringes or in single dose vials. They were also available in a multi-vial presentation, in which the antigens were supplied in separate vials from the nanoemulsion adjuvant, which should then be mixed together before use (*e*.*g*., Pandemrix and Arepanrix). These vaccines require storage at 2-8°C and must not be frozen. Unintentional exposure to freezing temperatures can lead to a significant damage to the vaccines. Unfortunately, it was estimated that 75-100% of vaccines are exposed to freezing temperatures in various segments of the supply chain [12]. Converting vaccines from liquid to dry powders is a promising approach to enhance their freezing and thermal stability. Freeze-drying is commonly used for the development of dry powder formulations of biologics including vaccines [13, 14]. Freeze-drying of vaccines containing O/W nanoemulsions is challenging, however, due to the sensitivity of nanoemulsions to the freezing and drying stress. Freezing of emulsions can result in phase separation, while dehydration can lead to interactions of surfactant molecules adsorbed on the oil droplets, which in turn adversely affects the emulsion stability and adjuvanticity [15, 16]. Nonetheless, there is evidence that it is possible to freeze-dry certain vaccine candidates that contain nanoemulsions [17]. For example, GLA-SE is a vaccine adjuvant currently under development. GLA-SE is composed of glucopyranosyl lipid A (GLA, a Toll-like receptor 4 (TLR-4) agonist) in squalene oil nanoemulsion (SE) [18]. A tuberculosis vaccine candidate comprised of ID93 antigen (*i*.*e*., a recombinant fusion protein antigen consisting of four *Mycobacterium tuberculosis* proteins) and the GLA-SE adjuvant (*i*.*e*., GLA-SE/ID93) was freeze-dried using D-trehalose and/or other disaccharides [18]. The mean hydrodynamic particle size of the GLA-SE/ID93 was about 80 nm and was increased by ≥10 nm after subjected to shelf freeze-drying and reconstitution [18, 19]. Vaccines containing MedImmune emulsions comprised of squalene oil, monophosphoryl lipid A or its synthetic analogue (PHAD), and Tween 80 exhibited a mean particle size increase from 70-90 nm to ∼110 nm or larger after subjected to shelf freeze-drying [15]. Importantly, the immune responses induced by the freeze-dried vaccine candidates were not different from their liquid counterparts when evaluated in animal models [15, 18].

Unfortunately, GLA-SE and the MedImmune emulsions are different from MF59 and AS03 in composition and physical properties (*e*.*g*., mean droplet size). Because the composition of nanoemulsion adjuvants is among the key factors that determine their susceptibility to drying [15], it remains unknown whether MF59, AS03, or nanoemulsions with the same or similar composition or droplet size as MF59 or AS03 and vaccines containing them can be converted to dry powders. The unique composition and physical properties of MF59 and AS03 may render them and vaccines containing them more sensitive to freezing and drying. For example, it was reported that nanoemulsions with smaller mean droplet size (*i*.*e*., ∼80 nm) can be easily freeze-dried as compared to nanoemulsions with relatively larger mean droplet size (*i*.*e*., 100-200 nm) [9]. The mean droplet size of GLA-SE and the MedImmune emulsions is ∼80 nm, but the mean droplet size of MF59 and AS03 is ∼160 nm.

The present study was designed to test the feasibility of applying TFFD to develop dry powders of AddaVax as well as vaccines containing AddaVax or MF59. AddaVax is preclinical grade nanoemulsion vaccine adjuvant with the same composition as MF59 (*i*.*e*., squalene oil (5% *v/v* or 4.29% *w/v* based on squalene density of 0.858 (PubChem CID: 638072)), Tween 80 (0.5% *w/v*) and Span 85 (0.5% *w/v*) in citrate buffer (10 mM, pH 6.5)) [20, 21]. AddaVax is for research use only. TFFD has been applied to successfully convert biologics, including vaccines, from liquid into stable dry powders [22-27]. It was hypothesized that the ultra-rapid freezing, relatively small gas-liquid interface, and low shear stress associated with the thin film freezing (TFF) process would minimize nanoemulsion droplet aggregation or fusion during the freezing step. We first tested the feasibility of thin-film freeze-drying AddaVax alone and then used ovalbumin (OVA) and lysozyme as model antigens to study the effect of stabilizing agent, antigen, buffer molarity, as well as TFF temperature on the particle size distribution of vaccines after subjected to TFFD and reconstitution. Finally, we applied TFFD to the Fluad Quadrivalent influenza virus vaccine that contains MF59 as well as mono-, bi-, and tri-valent (universal) influenza virus vaccine candidates containing H1 and/or H3 recombinant hemagglutinin (rHA) proteins and AddaVax. Although seasonal influenza occurs annually and seasonal flu vaccines are manufactured every year, the world is concerned about the threat of influenza pandemics [28], and there is an interest in developing universal flu vaccines. Our bivalent and trivalent influenza virus vaccine candidates can be considered universal flu vaccines. For universal flu vaccines, a shelf-life beyond the 6-9 months for seasonal flu vaccines is likely needed, and this need may be met by developing the vaccines into dry powders.

## 2. Materials and Methods

### 2.1 Influenza HA antigens and viruses

Two H3 (*i*.*e*., TJ-5 and J-4) and one H1 (*i*.*e*., Y-2) influenza virus rHA proteins were constructed using computationally optimized broadly reactive antigen (COBRA) methodology, a multiple-layered consensus building approach to design novel immunogens as vaccine candidates [29, 30]. The rHA proteins have a molecular weight of ∼270 kDa and were expressed in HEK-293 cells and purified using IMAC Sepharose^®^ High Performance resin (Sigma Aldrich, St. Louis, MO). Viruses were from the Influenza Reagents Resource (IRR) (Manassas, VA), BEI Resources (Manassas, VA), the Centers for Disease Control and Prevention (Atlanta, GA), or VIRAPUR, LLC (San Diego, CA, USA). Viruses were passaged once in the same growth conditions as they were received, in either embryonated chicken eggs or semi-confluent Madin-Darby Canine Kidney (MDCK) cell culture. H1N1 viruses used include A/Brisbane/02/2018, A/Michigan/45/2015, A/California/07/2009, A/Brisbane/59/2007, A/Solomon Islands/3/2006. H3N2 viruses used include A/South Australia/34/2019, A/Switzerland/8060/2017, A/Texas/71/2017, A/Kansas/14/2017, A/Singapore-IFNIMH-16-0019/2016, A/Hong Kong/4801/2014, A/Switzerland/9715293/2013, and A/Texas/50/2012.

### 2.2 Preparation of AddaVax dry powders by TFFD and conventional shelf freeze-drying

AddaVax (50 *μ*L, InvivoGen, San Diego, CA) was mixed with trehalose (Sigma-Aldrich) in citrate buffer (10 mM, pH 6.5) at a final sugar concentration of 30 mg/mL. The liquid AddaVax formulation was converted to a dry powder using either TFFD or conventional shelf freeze-drying. Thin-film freeze-dried powder was prepared as previously described [25]. Briefly, the liquid formulation (100 *μ*L) was dropped onto a cryogenically cooled stainless-steel surface (*i*.*e*., cylindrical drum) having a temperature of -100 °C. The liquid droplets were rapidly spread and frozen to form thin films. Shelf freeze-dried powder was prepared by gradually cooling 0.5 mL of the liquid AddaVax formulation containing trehalose at a concentration of 30 mg/mL from room temperature (∼ 21 ± 2°C) to -40°C at a cooling rate of about 2°C/min. The frozen liquid was maintained at -40°C for 1 h in the lyophilizer prior to lyophilization. Frozen thin-films prepared by TFFD and frozen liquid prepared by conventional shelf freezing were subsequently lyophilized using an SP VirTis AdVantage Pro Freeze Dryer with Intellitronics Controller (SP Scientific, Stone Ridge, NY). Lyophilization was performed over about 50 h at pressures ≤ 100 mTorr. The shelf temperature was maintained at -35°C for 30 h and then gradually ramped to +20°C throughout ∼16 h. During the secondary drying phase, the vials were kept at +20°C for 4 h. Vials were stoppered in nitrogen gas at 100 mTorr, sealed using aluminum caps, and then stored in a vacuum desiccator at room temperature until analysis. Reconstitution was performed by adding 100 *μ*L of milli-Q water. Reconstituted AddaVax formulations were diluted 20-fold with milli-Q water before droplet size measurements. Z-average hydrodynamic droplet size distribution was determined by dynamic light scattering (DLS) using a Malvern Zeta Sizer Nano ZS (Worcestershire, UK).

### 2.3 Effect of antigen, stabilizing agent, buffer molarity, freezing method, and TFF temperature on the particle size distribution of AddaVax-adjuvanted vaccines using OVA and lysozyme as model antigens

AddaVax-adjuvanted OVA model vaccine (AddaVax/OVA) formulations containing AddaVax (50 *μ*L), OVA (6 *μ*g, Sigma-Aldrich) and a stabilizing agent selected from sucrose (Merck KGaA, Darmstadt, Germany), D-mannitol and D-trehalose dihydrate (Sigma-Aldrich) at a concentration between 50 and 500 mg/mL were prepared by simple mixing (**Table 1**). Vaccine formulations (100 *μ*L) were subjected to TFFD as described above and hydrodynamic particle size distribution in the formulations was measured after reconstitution and dilution with milli-Q water. All formulations were in citrate buffer (pH 6.5) and frozen into thin-films at -100°C.

**Table 1.** Compositions of vaccine formulations prepared to investigate the effect of stabilizing agent and stabilizing agent concentration on the particle size distribution of AddaVax/OVA vaccine subjected to TFFD and reconstitution.

| Stabilizing agent |  |  | Citrate buffer molarity (mM) |
| --- | --- | --- | --- |
| Sugar/sugar alcohol | Concentration (mg/mL) | Mass fraction |  |
| Sucrose | 50 | 0.51 | 2.5 |
| Sucrose | 100 | 0.67 |  |
| Mannitol | 50 | 0.51 |  |
| Mannitol | 100 | 0.67 |  |
| Trehalose | 100 | 0.67 |  |
| Trehalose | 50 | 0.44 | 5 |
| Trehalose | 100 | 0.61 |  |
| Trehalose | 125 | 0.66 |  |
| Trehalose | 250 | 0.80 |  |
| Trehalose | 500 | 0.87 |  |

To study the effect of drying technology, freezing rate and repeated freezing, and thawing on the particle size of AddaVax/OVA vaccine, liquid formulation of AddaVax/OVA vaccine having the same composition of that subjected to TFFD (*i*.*e*., AddaVax (50 *μ*L), OVA (6 *μ*g) and trehalose (125 mg/mL in citrate buffer (2.5 mM, pH 6.5)) was converted to dry powder using shelf freeze-drying as described above. The effect of TFF and shelf freezing on mean particle size of AddaVax/OVA vaccine was also investigated. Frozen thin-films prepared by dropping 100 *μ*L of liquid vaccine formulation on a cryogenically cooled drum having a temperature of -100°C and frozen formulation prepared by shelf freezing (*i*.*e*., cooling 0.5 mL of the liquid vaccine formulation from room temperature (∼ 21 ± 2°C) to -40°C at a cooling rate of ∼ -2°C/min) were thaw at 4°C for 1 h. Formulations were adequately diluted in milli-Q water before determining their Z-average hydrodynamic particle size by DLS. The effect of repeated freezing and thawing on the mean particle size of liquid AddaVax/OVA formulation and its thin-film freeze-dried powder counterpart was also investigated. Liquid formulation (0.5 mL) and dry powder were subjected to three consecutive cycles of freezing at -20 °C for 8 h and thawing at 4 °C for 16 h. At the end of the third cycle, the powder was reconstituted and adequately diluted with milli-Q water and Z-average hydrodynamic particle size was determined by DLS.

Table 2 shows the composition of various vaccine formulations prepared to investigate the effect of TFF temperature (*i*.*e*., the temperature of the cryogenically cooled drum surface), the antigen, antigen amount and buffer molarity on the Z-average hydrodynamic particle size of AddaVax/OVA vaccine after subjected to TFFD and reconstitution. All formulations comprised 50 *μ*L of AddaVax and trehalose at a concentration of 125 mg/mL and were frozen into thin-films at a drum temperature of - 100°C. Relevant samples (**Table 2**) were also frozen into thin-films at drum temperatures of -50°C and -180°C to study the influence of TFF temperature on the Z-average hydrodynamic particle size of the model vaccine. Frozen vaccine thin-films were lyophilized and the particle size distribution upon reconstitution was determined as described above.

**Table 2.**
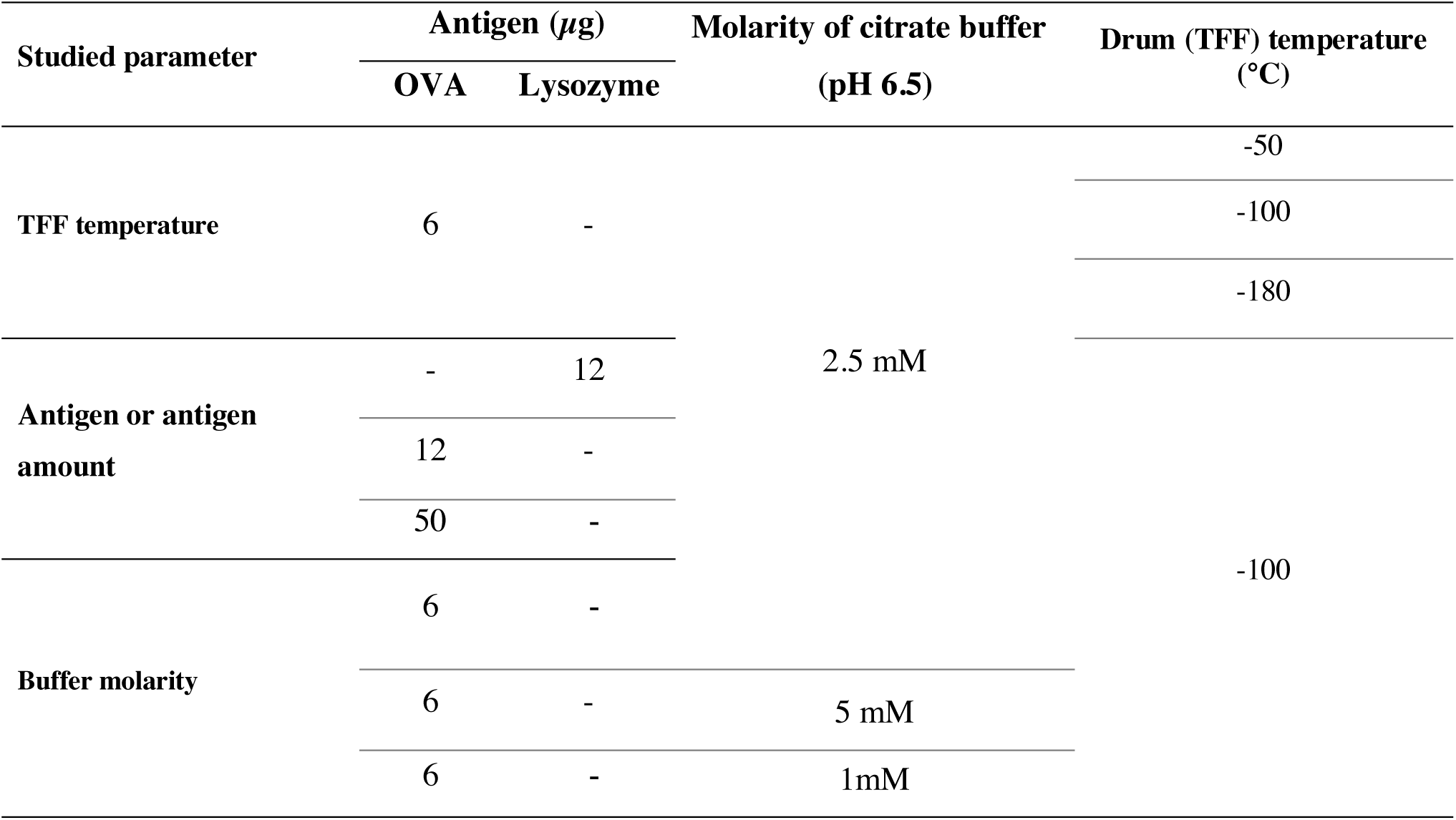
Compositions of vaccine formulations prepared to investigate the effect of TFF temperature, antigen, antigen amount and buffer molarity on particle size distribution of AddaVax-adjuvanted model vaccines subjected to TFFD and reconstitution.

### 2.4 Characterization of AddaVax/OVA dry powders

Thermal analysis of AddaVax/OVA powder prepared using the TFFD technology was performed using a differential scanning calorimeter Model Q20 (TA Instruments Inc., New Castle, DE) equipped with a refrigerated cooling system (RCS40, TA Instruments Inc.). Samples were first cooled down to -40°C at a ramp rate of 10°C/min, then ramped from -40°C to 300°C at a heating ramp rate of 5°C/min. Data were processed by TA Instruments Trios V.5.1.1.46572 software. Powder crystallinity was evaluated using a Rigaku Oxford Diffraction HyPix6000E Dual Source diffractometer (Tokyo, Japan) using a μ-focus sealed tube Cu Kα radiation source (λ = 1.5418Å) with collimating mirror monochromators. The instrument was operated at an accelerating voltage of 50 kV at 0.8 mA. The data were collected at 100 K using an Oxford Cryostream low temperature device (Oxford Cryosystems Ltd, Oxford, United Kingdom). A continuous ϕ rotation of the sample was maintained for each of three different orientations of the sample for 100 seconds for each frame. The three frames collected on the 2-dimensional detector were combined to generate a 1-dimensional powder pattern. The data collection and data reduction were performed using Rigaku Oxford Diffraction’s CrysAlisPro V 1.171.42.25a. Residual water content in the vaccine powder was quantified by volumetric Karl Fischer titration using a Mettler Toledo V20 titrator (Mettler Toledo, Columbus, OH).

### 2.5 Preparation of AddaVax-adjuvanted influenza vaccine powders and Fluad Quadrivalent dry powder using TFFD

Four influenza vaccine formulations were prepared by mixing one or more rHA proteins at a total amount of 6 *μ*g with 50 *μ*L of AddaVax (**Table 3**). Trehalose in citrate buffer (2.5 mM, pH 6.5) was employed as a stabilizing agent at a concentration of 125 mg/mL. Formulations (100 *μ*L) were thin-film frozen at -100°C and lyophilized as described above. Fluad Quadrivalent vaccine was donated by HEB Pharmacy with approval of the Texas State Board of Pharmacy. Fluad Quadrivalent vaccine contains MF59 and the HA proteins of four influenza strains at 15 *μ*g per 0.5 mL each. Trehalose dissolved in citrate buffer (2.5 mM, pH 6.5) (50 *μ*L) was mixed with 50 *μ*L of the Fluad Quadrivalent vaccine to reach a final trehalose concentration of 125 mg/mL. The resultant liquid Fluad Quadrivalent vaccine formulation was then frozen to thin-films at -100°C and dried as described above.

**Table 3.** Compositions of AddaVax-adjuvanted monovalent, bivalent, and trivalent influenza virus vaccines.

| Vaccine candidate | AddaVax<br>($\mu\text{L}$) | rHA antigen(s) ($\mu\text{g}$) | | |
| --- | --- | --- | --- | --- |
|  |  | Y-2 | J-4 | TJ-5 |
| AddaVax/Y-2 | 50 | 6 | - | - |
| AddaVax/J-4 | 50 | - | 6 | - |
| AddaVax/Y-2/J-4 | 50 | 3 | 3 | - |
| AddaVax/Y-2/J-4/TJ-5 | 50 | 2 | 2 | 2 |

### 2.6 Characterization of AddaVax-adjuvanted influenza vaccine and Fluad Quadrivalent vaccine powders

Thin-film freeze-dried vaccine powders were reconstituted in milli-Q water, and particle size distribution, polydispersity index (PDI) and zeta potential values were meausred using a Malvern Zeta Sizer Nano ZS after dilution with milli-Q water. The integrity of HA proteins was investigated using SDS-PAGE analysis. Samples for SDS-PAGE analysis and hemagglutination assay were reconstituted in 50 *μ*L milli-Q water so that the HA content is 6 *μ*g/50 *μ*L to facilitate the analysis. Briefly, 10 *μ*L of reconstituted representative influenza vaccine (*i*.*e*., AddaVax/Y-2) of Fluad Quadrivalent was mixed with Laemmli Sample Buffer (Bio-Rad, Hercules, CA) and β-mercaptoethanol (2%, *v/v*, Sigma-Aldrich). Samples were heated at 95°C for 5 min prior to loading onto 4-20% Mini-PROTEAN^®^ TGX™precast polyacrylamide gel (Bio-Rad). Gel electrophoresis was done at 100 V for 90 min. Gel was stained in a Bio-Safe™ Coomassie G-250 Stain (Bio-Rad).

The integrity of the HA proteins in the TFFD powders was evaluated using a standard hemagglutination assay using chicken red blood erythrocytes as previously described [31]. Briefly, vaccine powders were reconstituted in water, and then 50 *μ*L sample was 2-fold serially diluted using phosphate buffered saline (PBS, 10 mM, pH 7.2) in U-bottom 96-well plates. The samples were then incubated with 50 *μ*L of 1% chicken erythrocyte suspension (Rockland Immunochemicals, Inc., Limerick, PA) in PBS at room temperature for 30 min. Hemagglutination titers were reported as the reciprocal of the last dilution where hemagglutination was observed (*i*.*e*., absence of chicken erythrocyte precipitation) and were expressed in hemagglutination units (HAUs)/50 *μ*L.

### 2.7 Animal studies

Female BALB/c mice (6 to 8 weeks old) were from Jackon Laboratory (Bar Harbor, ME) and housed in microisolator units. The mice were allowed free access to food and water and were cared for under USDA Guidelines for Laboratory Animals. All procedures were reviewed and approved by the Institutional Animal Care and Use Committee at the University of Georgia. Mice (10 per group) were intramusculary injected twice, four-weeks apart, with different influenza virus vaccine formulations at a dose that contained 3 *μ*g rHA protein(s) per mouse (**Table 4)**. Mice were bled in weeks 0, 4 and 8. For viral challenge, mice were briefly anesthetized and infected with 50 *μ*L A/Kansas/14/2017 H3N2 intranasally (5 × 10^6^ PFU). Mice were monitored for weight loss and euthanized 14 days after challenge. Weight loss more than 25% was used as a primary measurement for determination of humane endpoint. Also, dyspnea, lethargy, response to external stimuli and other respiratory distress was closely monitored for the determination of humane endpoint.

**Table 4.**
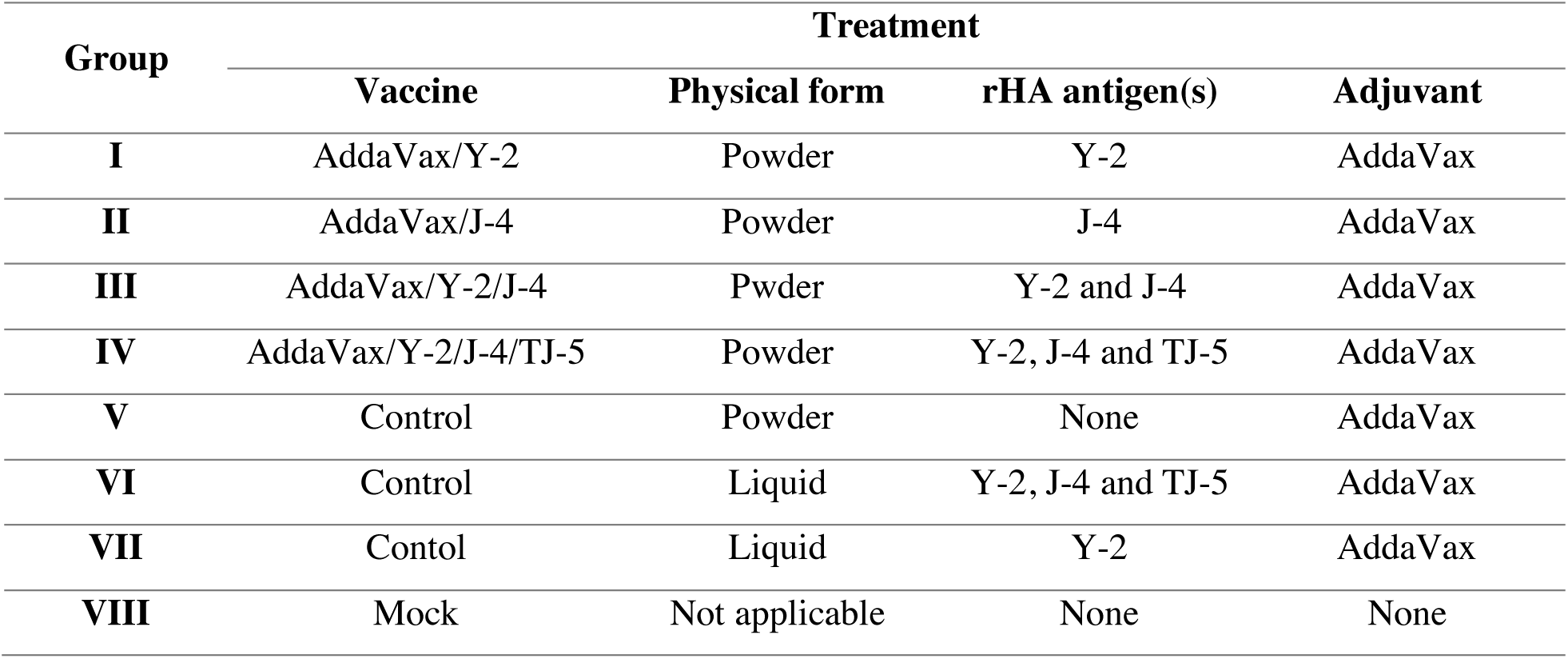
Animal study design.

| Group | Treatment |  |  |  |
| --- | --- | --- | --- | --- |
|  | Vaccine | Physical form | rHA antigen(s) | Adjuvant |
| I | AddaVax/Y-2 | Powder | Y-2 | AddaVax |
| II | AddaVax/J-4 | Powder | J-4 | AddaVax |
| III | AddaVax/Y-2/J-4 | Powder | Y-2 and J-4 | AddaVax |
| IV | AddaVax/Y-2/J-4/TJ-5 | Powder | Y-2, J-4 and TJ-5 | AddaVax |
| V | Control | Powder | None | AddaVax |
| VI | Control | Liquid | Y-2, J-4 and TJ-5 | AddaVax |
| VII | Control | Liquid | Y-2 | AddaVax |
| VIII | Mock | Not applicable | None | None |

### 2.8 Hemagglutination inhibition assay

The hemagglutination inhibition (HAI) assay was used to assess the ability of anti-rHA protein antibodies to inhibit hemagluttination of erythrocytes by a panel of H1N1 and H3N2 viruses. The protocols were adapted from the WHO laboratory influenza surveillance manual [32]. Briefly, sera from mice 4 weeks after boost immunization were treated with receptor-destroying enzyme (RDE) (Denka Seiken, Japan) to inactivate nonspecific inhibitors. RDE was added 3:1, *w/v*, to sera and incubated 18 h at 37°C. RDE was then inactivated by incubation at 56°C for 45 min. RDE-treated sera were brought up to a final 1:10 mixture in PBS and then diluted in 2-fold serially in V-bottom microtiter plates. An equal volume of virus, adjusted to 8 HAUs/50 *μ*L, was added to each well. H3N2 virus was adjusted with 20 nM Oseltamivir. After 20 min incubation, 0.8% erythrocytes in PBS were added (Lampire Biologicals, Piperville, PA). For H1N1, turkey erythrocytes were used. For H3N2, guinea pig erythrocytes in precence 20 nM Oseltamivir were used.

### 2.9 Focus reduction assay

Focus Reduction Assay (FRA) was used to assess the ability of polyclonal sera from vaccinated mice to neutralize H1N1 and H3N2 viruses *in vitro* as previously described [33]. Briefly, MDCK-SIAT1 cells were plated at 5 × 10^4^ cells per well in a 96-well plate in media (DMEM containing 5% heat-inactivated fetal bovine serum and penicillin-streptomycin). RDE-treated mouse sera were serially diluted 2-fold starting at 1:20 dilution in virus growth medium (DMEM containing 0.1% BSA, penicillin-streptomycin, and 1 *μ*g/mL TPCK-treated trypsin). Sera (50 *μ*L) was added to the cell monolayers. Afterward, 50 *μ*L of virus (600 focus forming units (FFU)/50 *μ*L) were added and the plates were incubated for 2 h at 37°C with 5% CO_2_. The cells were overlaid with 1.2% Avicel (FMC Health and Nutrition, Philidelphia, PA) in 2× modified Eagle medium containing 0.1% BSA, penicillin-streptomycin, and 1 *μ*g/mL TPCK-treated trypsin. Plates were incubated for 24 h at 37°C with 5% CO_2_.

The overlays were removed and the cell monolayers washed with PBS to remove any residual Avicel. The plates were fixed with 4% formalin and cells were permeablized with 0.5% Triton X-100 in PBS/glycine. The plates were washed with PBS containing 0.1% Tween 20 and incubated for 1 h at room temperature with a monoclonal antibody against influenza virus A or B nucleoprotein from The IRR. After washing three times, the cells were incubated with goat anti-mouse peroxidase labeled IgG (474-1802; SeraCare, Inc, Milford, MA) for 1 h at room temperature. The plates were washed three times and infectious foci were visualized by adding TrueBlue substrate (SeraCare) containing 0.03% H_2_O_2_ to the cells for 10 min at room temperature. The reaction was stopped by washing with distilled water five times. The foci were counted using a BioSpot analyzer with ImmunoCapture 6.4.87 software (CTL, Shaker Heights, OH). The virus control well containing no sera was used for comparison of focus reduction.

### 2.10 Statistical analysis

Student’s t-test or One-way ANOVA followed by Tukey’s or Dunnet multiple comparison test were performed using GraphPad Prism version 8.0.0 for Windows (GraphPad Software, San Diego, CA). Differences were deemed significant if *p ≤ 0*.*05*.

## 3. Results and discussion

### 3.1. Thin-film freeze-drying of AddaVax adjuvant

O/W nanoemulsions such as MF59 are vaccine adjuvants that can elicit robust antibody and cellular immune responses. O/W nanoemulsion-adjuvanted vaccines are freeze-sensitive. Formulating these vaccines as dry powders can potentially address their freezing sensitivity. Initially, we tested the applicability of TFFD technology for converting AddaVax, a preclincial grade equivalent of MF59, into a dry powder without adversely affecting its droplet size and size distribution. Our previous work showed that trehalose at low concentration can protect lipid-based vaccine adjuvants (*e*.*g*., liposomes) against freezing and/or drying-induced particle aggregation (unpublished data). Thus, liquid AddaVax formulation containing trehalose at a concentration of 30 mg/mL in citrate buffer (10 mM, pH 6.5) was frozen into thin films at a temperature of -100ºC followed by lyophilization. The same formulation was also converted to dry powder by standard shelf freeze-drying as a control. Approved seasonal and pandemic influenza vaccines comprise either AS03 or MF59 adjuvant equivalent to 10.69 mg or 9.75 mg of squalene oil, respectively, per 0.5 mL dose. In this study, the formulation contained 50 *μ*L of AddaVax per 100 *μ*L, which is equivalent to 10.7 mg squalene oil per 0.5 mL.

Dry powders of AddaVax prepared by TFFD or conventional shelf freeze-drying were reconstituted in water and mean droplet size of the nanoemulsion was determined by DLS. The droplet size is among the key quality attributes of nanoemulsion vaccine adjuvants [15]. It can affect both the nanoemulsion’s stability and adjuvanticity [15]. Thus, maintaining the integrity and size uniformity of nanoemulsion droplets in the dry powders is critical. As depicted in **Figure 1A**, the mean droplet size of AddaVax nanoemulsion did not significantly changed after it was converted to a dry powder using TFFD and reconstituted in water. On the contrary, shelf freeze-drying had a deleterious effect on the particle size distribution of AddaVax nanoemulsion, leading to significant particle aggregation or fusion as demonstrated by the extra groups of particulates or droplets in the range of 1 *μ*m and 5 *μ*m (**Figure 1B**). Thus, the Z-average mean droplet size of AddaVax in the dry powder prepared using shelf freeze-drying was significantly increased. Since the frozen thin-films prepared by TFF and the frozen liquid prepared by shelf freezing were dried using the same lyophilization cycle, the observed different effects of these dry powder engineering technologies on the nanoemulsion droplet size can be attributed to the freezing step.

**Figure 1.**
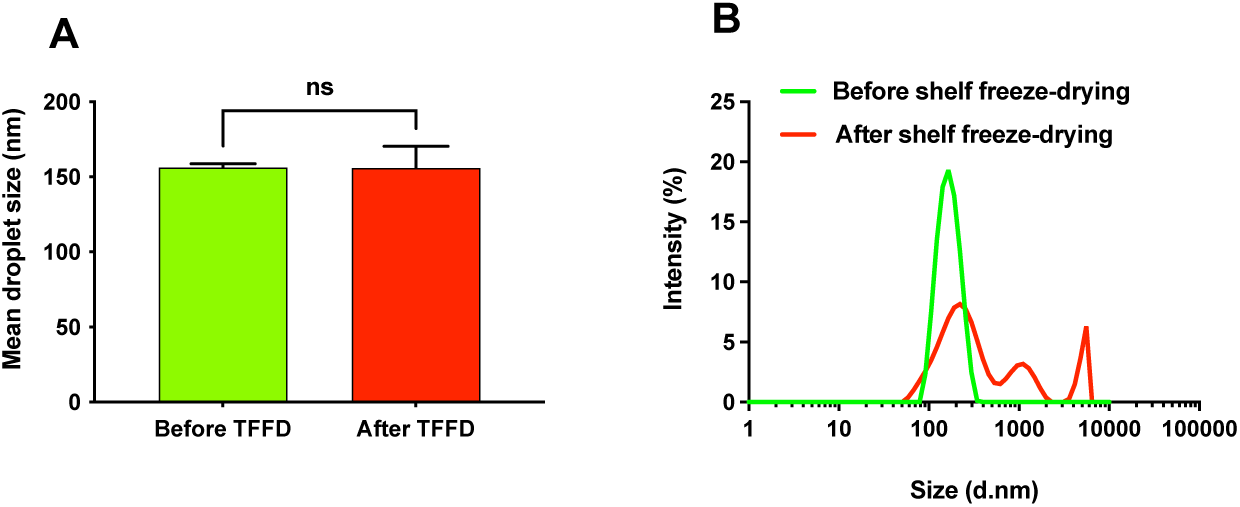
Effect of TFFD and shelf freeze-drying on AddaVax droplet size. (A) Mean droplet size of AddaVax after TFFD and reconstitution in water as compared to the liquid adjuvant (*i*.*e*., before TFFD). (B) Droplet size distribution of reconstituted AddaVax dry powder prepared using conventional shelf freeze-drying as compared to the liquid AddaVax before subjected to shelf freeze-drying. *ns: non-significant (p>0*.*05)*.

### 3.2. Thin-film freeze-drying of AddaVax-adjuvanted vaccines using OVA or lysozyme as model antigens

#### 3.2.1. Effect of stabilizers

AddaVax was successfully converted to dry powder using TFFD technology without a significant effect on mean droplet size and size distribution of the nanoemulsion after reconstitution. Then, the applicability of TFFD for converting AddaVax-adjuvanted vaccines into dry powders was investigated using sucrose, trehalose or mannitol as a stabilizing agent at a concentration of 100 mg/mL and OVA (6 *μ*g) as a model antigen. Appropriate stabilizing excipient(s) must be incorporated in the formulation in order to protect the nanoemulsion droplets against possible freezing and/or the drying-induced stresses [14, 22]. All excipients have helped to maintain AddaVax/OVA vaccine’s monodispersed particle size distribution after TFFD and reconstitution (**Figure 1S**); however, the vaccine’s mean particle size was increased (**Figure 2A**). Sucrose was more effective in maintaining the vaccine mean particle size after TFFD than trehalose (*p<0*.*05*) or mannitol (*p<0*.*0001*). Sucrose as a stabilizing excipient has resulted in a particle size increase by 42 ± 9 nm (*i*.*e*., 26%) after TFFD. When trehalose or mannitol was used as a stabilizing excipient at the same concentration level, the mean particle size increased by 63 ± 6 nm or 101 ± 4, respectively. This is in agrement with a previous report that sucrose was more effective than trehalose and mannitol in preserving the particle size of GLA-SE/ID93 vaccine after freeze-drying [19].

**Figure 2.**
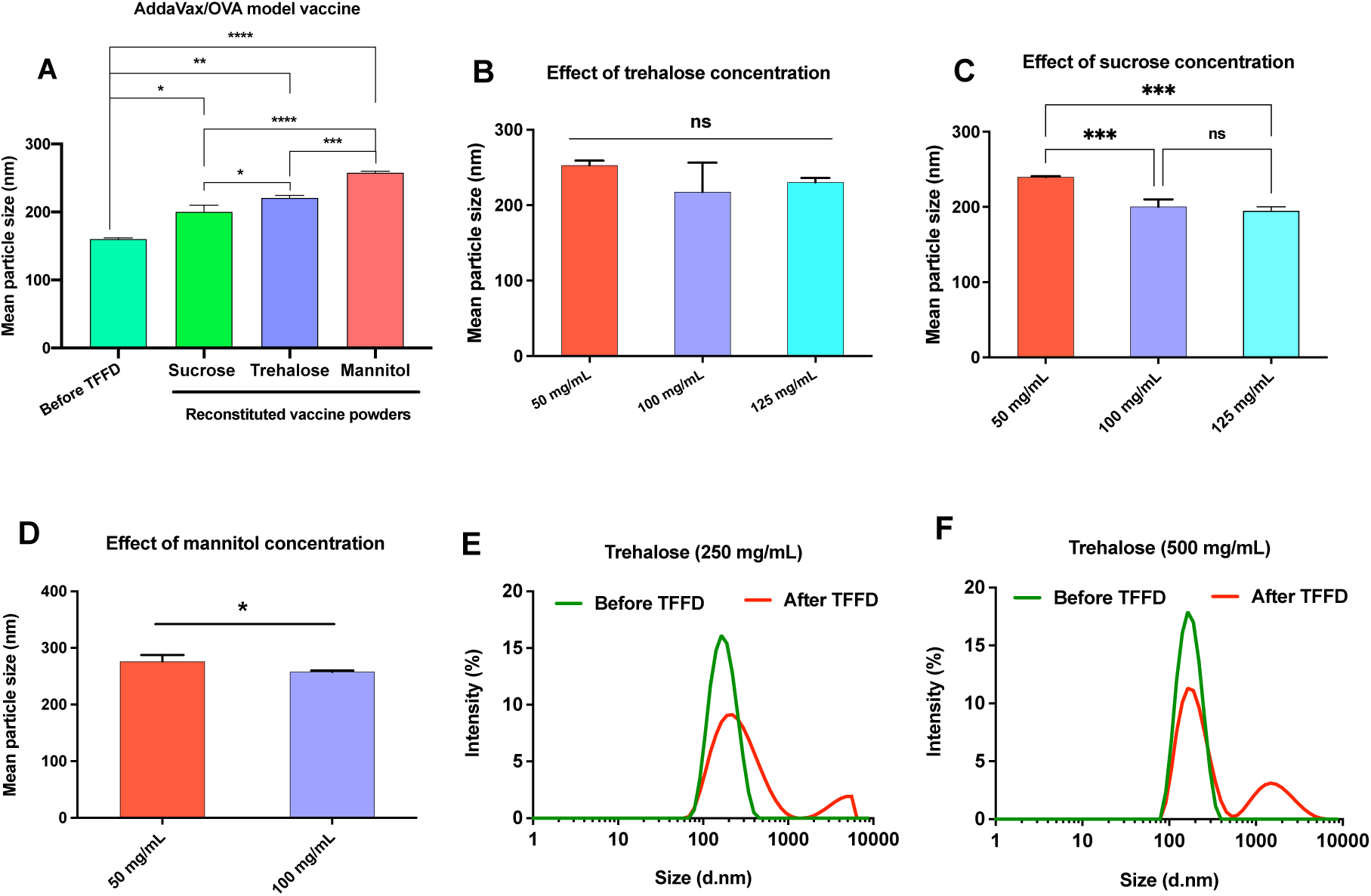
Effect of stabilizing agent and its concentration on mean particle size of AddaVax/OVA model vaccine. (A) The effeciency of sucrose, trehalose and mannitol in terms of maintaining the vaccine mean particle size after TFFD and reconstitution was invesitgated at sugar/sugar alcohol concentration of 100 mg/mL. Liquid vaccine fromulations (*i*.*e*., before TFFD) comprising different stabilizing agents at 100 mg/mL showed similar mean droplet size. (B-F) Effect of stabilizing agent concentration on the mean particle size of AddaVax/OVA powder prepared using TFFD after reconstitution in water. *\*p<0*.*05*, \*\**p*<0.01, *\*\*\*p<0*.*001, ****p<0*.*0001, ns*: non-significant (*p>0*.*05*).

In addition to the identity of the stabilizing agent, its concentration in the formulation is also critical [34]. Thus, the stabilizing effect of various stabilizing agents at different concentrations on the vaccine particle size was investigated (**Figure 2B-F**). Overall, it appeared that sucrose or trehalose at 100 or 125 mg/mL was most effective in minimizing the increase in the particle size of the AddaVax/OVA vaccine upon TFFD and reconstitution. Mannitol was less effective, likely because mannitol crystallizes during freezing [35]. Trehalose at 100 or 125 mg/mL was also effective, but not as effective as sucrose. It was noted that trehalose at 250 mg/mL and 500 mg/mL resulted in bimodal particle size distribution in the reconstituted model vaccine (*i*.*e*., large aggregates with mean droplet size >1*μ*m were observed) (**Figure 2E-F**). Generally, the higher the concentration of the stabilizing agent, the better its stabilizing effect. However, dispersion destabilization can be induced when the excipient’s concentration required for optimal stability is exceeded [36].

Formulations containing various stabilizers at a concentration of 50 mg/mL were selected to investigate the crystallinity of their powders because they showed a clear distinction in their Z-average hydrodynamic particle size (**Figure 2B-D**). As depicted in **Figure 3A**, TFF of vaccine formulation containing β-D-mannitol at a concentration of 50 mg/mL resulted in the crystallization of β-D-mannitol mainly in the δ and α polymorphs with the δ polymorph dominating. Our previous work showed also that mannitol crystallizes during TFF in the α polymorph [24] or β and δ polymorphs [37]. The distribution of mannitol crystal forms depends on the formulation compostion as well as the freezing and drying conditions [38]. For instance, in this study β-D-mannitol solution in water (50 mg/mL) crystallized in β and α polymorphs (data not shown). On the other hand, AddaVax/OVA powders containing either sucrose or trehalose were amorphous (**Figure 3A**). To be effective in protecting nanoemulsions against freezing and/or drying stress, stabilizing agents must retain amorphous structures during the freezing and drying steps [39, 40]. For instance, crystalline lyophilisate was reported to increase the mean particle size of GLA-SE after reconstituion [19], possibly due to destabilization of the nanoemulsion’s membrane [41]. It is noteworthy that mannitol at low concentration (*i*.*e*., 0-1% *w/v*) promotes the formation of amorphous lyophilizate [19]; however, in this study mannitol at low concentrations was ineffective as a stabilizing agent. Consequently, mannitol was excluded from further investigations. Sucrose is a non-crystallizing sugar [42] that remains amorphous after TFFD and thus can maintain the integrity of the nanoemulaion’s droplets. Trehalose crystallizes as trehalose dihydrate during the freezing step, which in turn undergoes dehydration to amorphous anhydrate during the drying step [39]. Crystallization of trehalose during the freezing step can justify its relatively lower efficiency as a stabilizing excipient as compared to the non-crystallizing sucrose (**Figure 2**).

**Figure 3.**
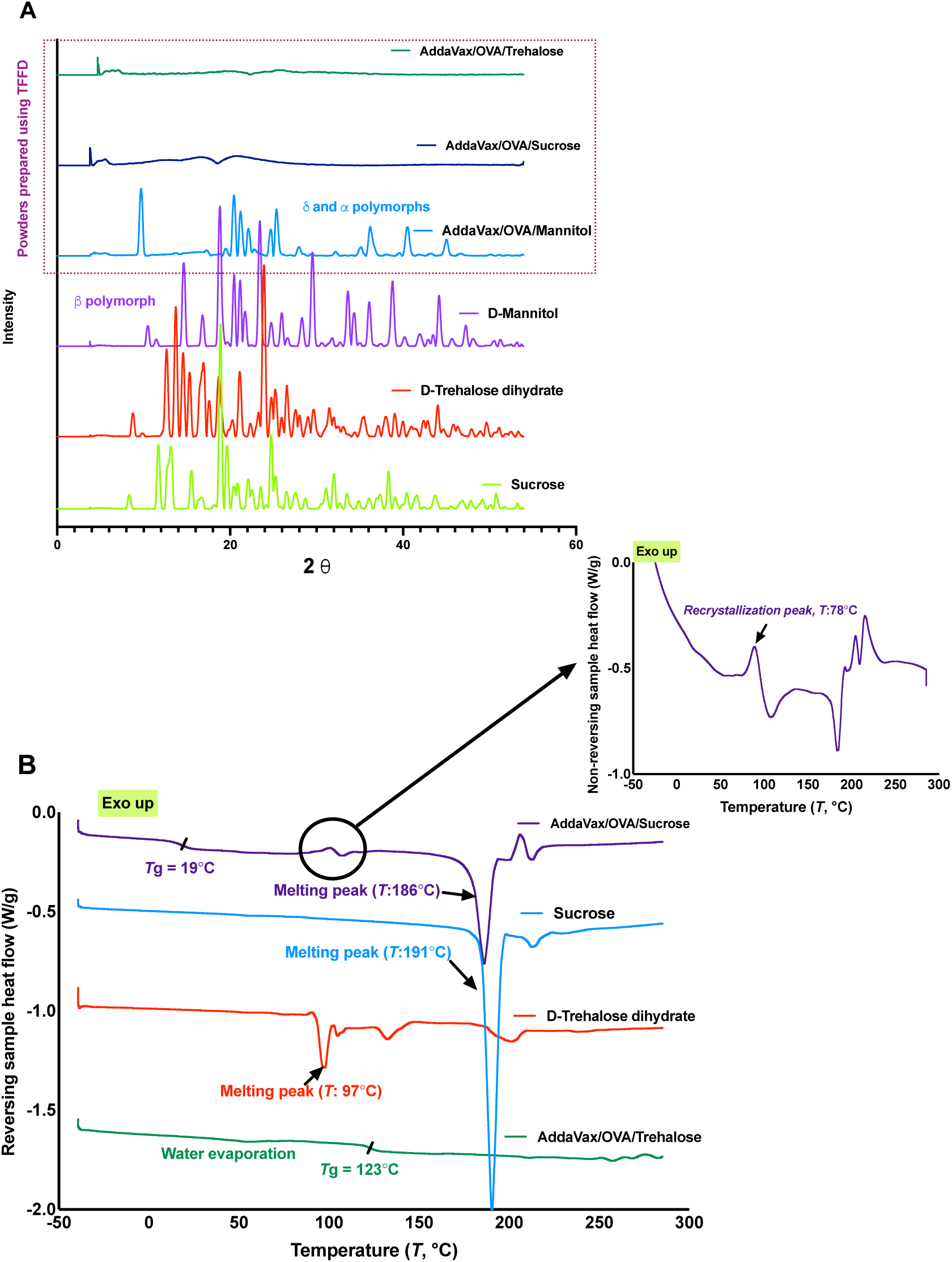
Characterization of AddaVax/OVA dry powders. (A) Powder X-ray diffraction (PXRD) patterns of thin-film freeze-dried AddaVax/OVA vaccine powders prepared with sucrose, trehalose or mannitol as a stabilizer as well as pure stabilizers as controls. (B) DSC thermograms of thin-film freeze-dried AddaVax/OVA vaccine powders prepared with trehalose (*i*.*e*., AddaVax/OVA/Trehalose) or sucrose (*i*.*e*., AddaVax/OVA/Sucrose) as a stabilizer and pure sugars as controls. Inset, the non-reversing heat flow thermogram of AddaVax/OVA/sucrose showing a recrystallization exothermic peak of sucrose.

As mentioned above, sucrose and trehalose were both effective when they were incorporated in the AddaVax/OVA formulation at a concertation of 100 mg/mL or 125 mg/mL. Sucrose is more hygroscopic than trehalose [43]. Water sorption by sucrose-based dry powders can deteriorate their physical properties [14, 22, 44]. As shown in **Figure 3B**, the glass transition temperature (*T*g) of AddaVax/ OVA model vaccine containing trehalose or sucrose as a stabilizing excipient was 123°C and 19°C, respectively. The observed *T*g of amorphous sucrose is lower than the reported values (*i*.*e*., 52-79°C) [45], which could be due to the relatively high residual water content in the vaccine dry powder (∼5%). Thus, trehalose can be more effective than sucrose in long-term stabilization of vaccine powder due to its high *T*g [40]. Furthermore, trehalose can slow the crystallization of low *T*g formulations to an extent that they can be stored at ambient temperatures [45]. The *Tg* of the developed vaccine powder comprising trehalose as a stabilizer is sufficiently higher than room temperature and thus, the powder has the potential to be stored in ambient temperatures [22]. Therefore, the vaccine formulation comprising AddaVax (50 *μ*L), OVA (6 *μ*g), and trehalose (125 mg/mL) in citrate buffer (2.5 mM, pH 6.5) was selected for further investigations unless otherwise described.

#### 3.2.2. Effect of freezing process (freezing rate) and repeated freezing and thawing on the particle size of AddaVax/OVA vaccine

Shelf freeze-drying had a deleterious effect on the particle size distribution of AddaVax/OVA vaccine and led to an extra group of particulates in the range of 1000 nm (**Figure 4A-B**). Although the dry powder of AddaVax/OVA vaccine prepared using TFFD maintained a unimodal particle size distribution after reconstitution (PDI = 0.26 ± 0.02), vaccine mean particle size increased from 159 ± 1 nm to 229 ± 10 nm (*p<0*.*05*). Since the freezing step is considered the most critical step for the integrity of freeze-dried emulsions [17, 46], the effect of freezing rate on the mean particle size of the vaccine was investigated. As shown in **Figure 4C-D**, the thin-film freezing step contributed to the increase of vaccine mean particle size to a smaller extent as compared to the the drying step (by 29 nm *vs* 41 nm, respectively, *p<0*.*05*). It was reported that coating of the oil droplet surface with trehalose through hydrogen bonding during the drying step can increase the hydrodynamic particle size by up to 80 nm [47]. The drying step was also responsible for the increase in the polydispersity index of the vaccine (**Figure 4E**). On the contrary, the deleterious effect of shelf freeze-drying process on the Z-average hydrodynamic partice size of the vaccine could be mainly attributed to the freezing step (**Figure 4F**). Nonetheless, the drying step had also lead to a slight increase of the vaccine hydrodynamic particle size (**Figure 4F**), likely in part due to coating of the particles in the vaccine by trehalose.

**Figure 4.**
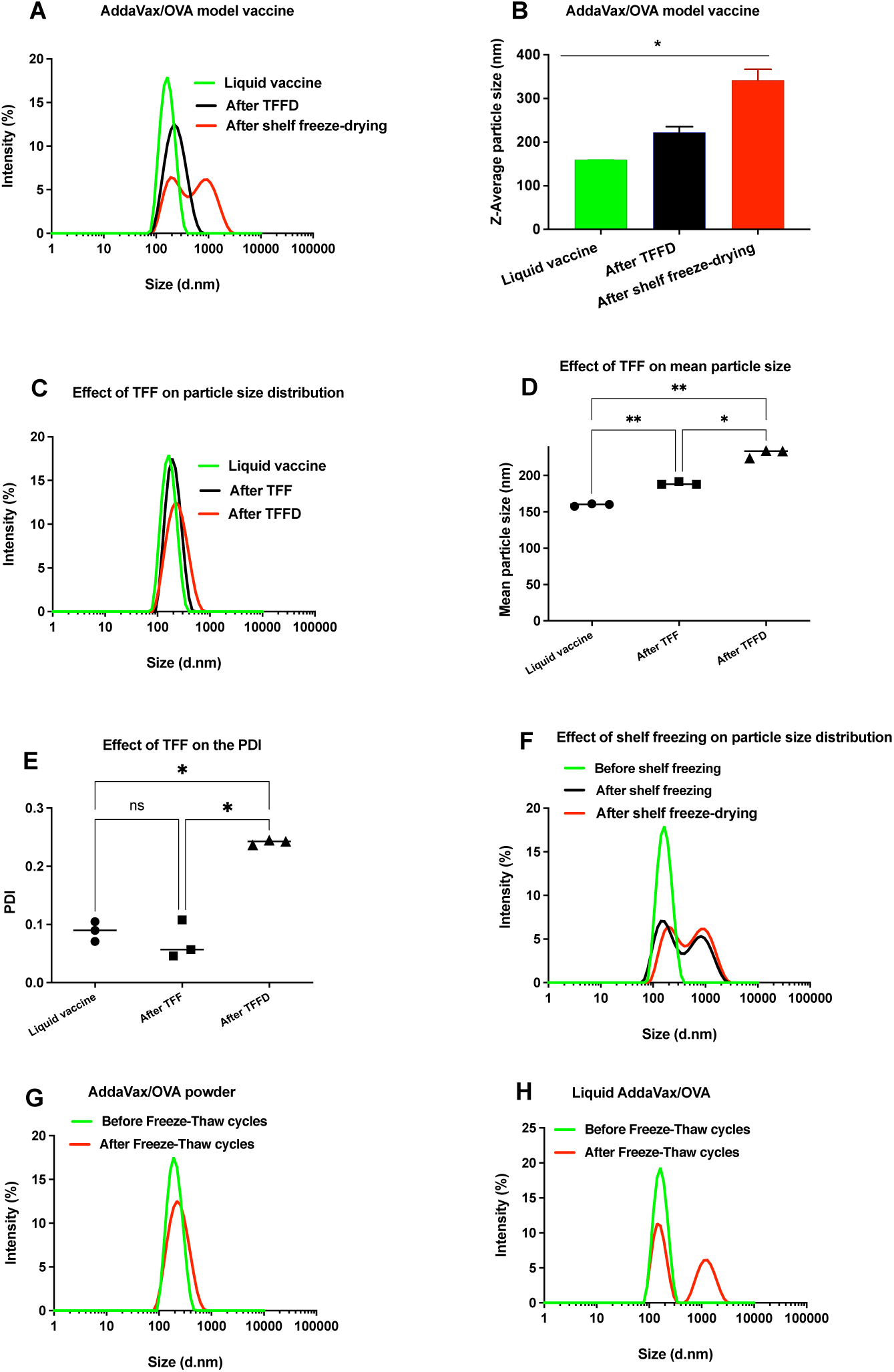
Effect of freezing process (freezing rate) and repeated freezing and thawing on the Z-average hydrodynamic particle size of AddaVax/OVA vaccine. (A) Particle size distribution and (B) Z-average hydrodynamic particle size of liquid and reconstituted AddaVax/OVA vaccine dry powders prepared using shelf freeze-drying or TFFD. (C-E) Effect of TFF on particle size distribution, Z-average hydrodynamic particle size and PDI of AddaVax/OVA vaccine. (F) Effect of shelf freezing on particle size distribution of AddaVax/OVA vaccine. (G-H) Effect of repeated freezing and thawing on intensity particle size distribution of AddaVax/OVA vaccine as a thin-film freeze-dried powder or in liquid. *\*p<0*.*05*, \*\**p*<0.01, *ns*: non-significant (*p>0*.*05*).

The formation of ice crystals during freezing of emulsions can induce the aggregation and/or fusion of emulsion droplets as a result of surfactant layer aggregation [17, 46]. Addtionally, slow freezing results in large ice crystals and hence larger supercooling effects than fast freezing [48, 49]. Thus, the slow shelf freezing may have resulted in the bimodal particle size distribution in the AddaVax/OVA powder prepared by shelf freeze-drying. Consequently, we hypothesised that powder engineering technologies that achieve sufficientlly rapid cooling rates can protect the nanoemulsion droplets against aggregation or fusion. TFF achieves cooling rates (*i*.*e*., 10^2^-10^3^ K/s) intermediate between spray freeze (*i*.*e*., 10^6^ K/s) and conventional shelf freezing (*i*.*e*., 0.017 K/s) [25]. The high freezing rates during TFF result in the formation of small ice crystals and homogenous distribution of the stabilizing agent [50]. Additionally, the large number of nuclei and thin ice channels formed during TFF prevent particle growth [26].

According to prescribing information, liquid nanoemulsion-adjuvanted vaccines (*e*.*g*., Fluad and Fluad Quadrivalent) should not be exposed to freezing. They should be discarded if thery were accidentally exposed to a freezing temperature which can be critical in case of pandemics. To test whether thin-film freeze-dried vaccines containing an nanoemulsion as an adjuvant is sensitive to freezing, thin-film freeze-dried AddaVax/OVA vaccine powder was subjected to three cycles of freezing and thawing to test its freezing sensitivity. As illustrated in **Figure 4G**, the mean particle size of the AddaVax/OVA vaccine dry powder was preserved after it was exposed to repeated freezing and thawing. On the contrary, repeated freezing and thawing of the liquid AddaVax/OVA vaccine resulted in significant particle aggregation **(Figure 4H)**. Freezing results in the formation of a network of crystalline oil droplets, and the network collapses and droplets undergo coalescence during thawing [51]. These results demonstrate the benefits of converting vaccines containing nanoemulsions into dry powders.

#### 3.2.3. Effect of TFF temperature, antigen, antigen amount and buffer molarity on the particle size of AddaVax/OVA vaccine

The applicability of TFFD for converting AddaVax-adjuvanted vaccines containing different antigens to dry powders was investigated using lysozyme or influenza virus rHA proteins (*i*.*e*., Y-2, J-4 and TJ-5). As shown in **Figure 5A-C**, the Z-average hydrodynamic particle size values of adjuvanted influenza vaccine candidates and lysozyme model vaccine after subjected to TFFD and subsequent reconstitution did not significantly increase (*p>0*.*05*), unlike the AddaVax/OVA vaccine, pointing out the effect of antigen on the particle size in thin-film freeze-dried AddaVax-adjuvanted vaccines. To test whether other factors may be adjusted to minimize the increase in the hydrodynamic particle size of the AddaVax/OVA vaccine after subjected to TFFD, TFF temperature, buffer molarity and antigen amount were explored. In addtion to affecting the freezing rate, the freezing temperature also affects the crystal growth [50]. Our results showed that the AddaVax/OVA vaccine thin films can be prepared at a wide range of temperatures without a significant effect on the mean particle size of the vaccine (**Figure 5D**). Moreover, the buffer molarity and antigen amount did not appear to have any effect on the Z-average hydrodynamic particle size of AddaVax/OVA vaccine after subjected to TFFD and subsequent reconstitution (**Figure 5E-F**). Therefore, the vaccine formulation comprising AddaVax (50 *μ*L), antigen (6 *μ*g), and trehalose (125 mg/mL) in citrate buffer (2.5 mM, pH 6.5) in 0.1 mL and thin-film frozen at -100 °C was used for additional studies.

**Figure 5.**
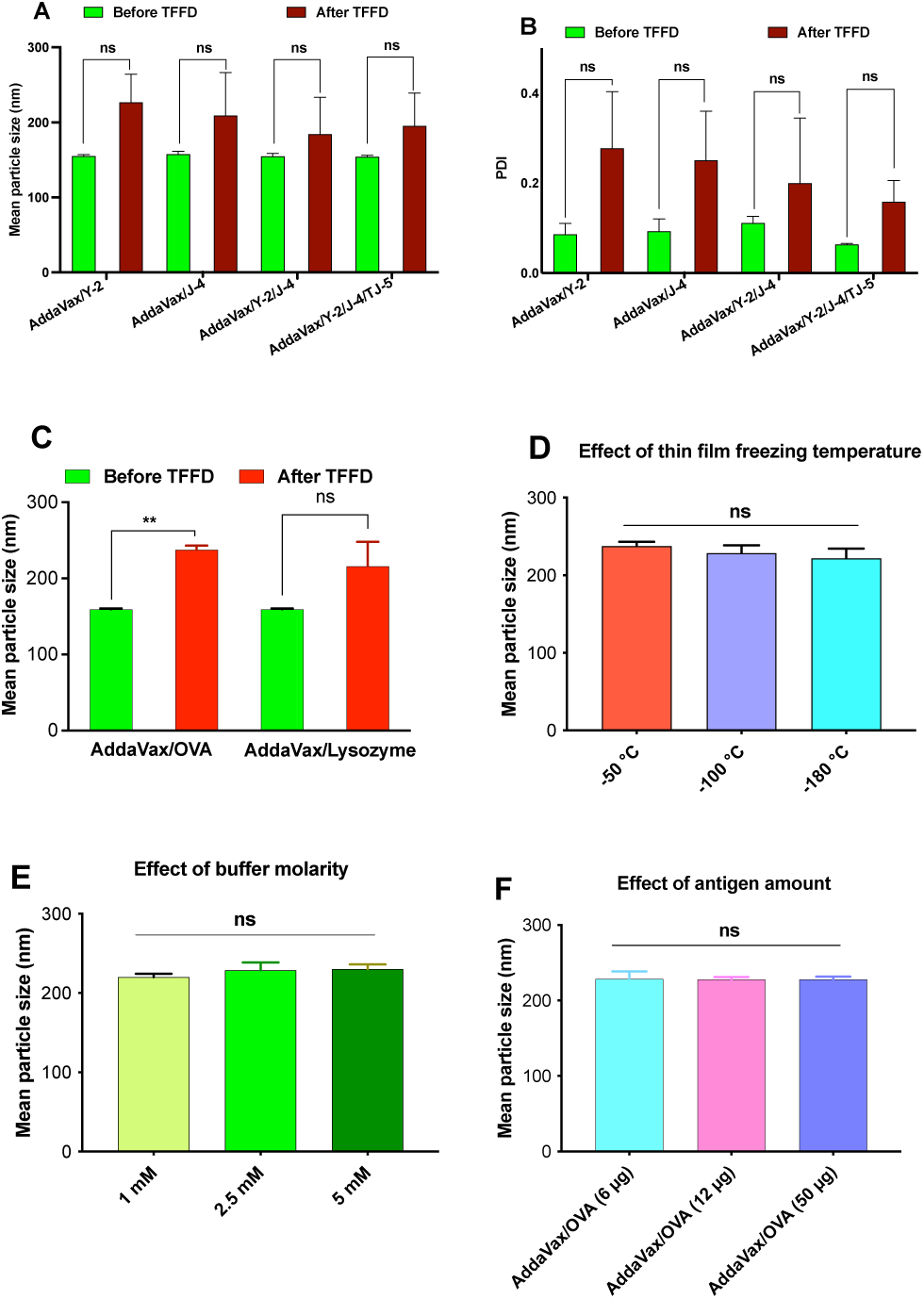
Effect of different antigens, antigen amount, buffer molarities, and thin-film freezing temperature on AddaVax-adjuvanted vaccines. (A) Z-average hydrodynamic particle size and (B) PDI values of AddaVax-adjuvanted influenza vaccines containing one or more rHA proteins at a total amount of 6 *μ*g/100 *μ*L. (C) Z-average hydrodynamic particle size of two model vaccines comprising 12 *μ*g/100 *μ*L of OVA (*i*.*e*., AddaVax/OVA) or lysozyme (*i*.*e*., AddaVax/lysozyme) as an antigen. (D) The effect of TFF temperature (*i*.*e*., drum temperature) on the Z-average hydrodynamic particle size of AddaVax/OVA comprising OVA (6 *μ*g/100 *μ*L) was studied at three drum temperatures (*i*.*e*., -50, -100, and -180°C). (E) The effect of buffer molarity on the Z-average hydrodynamic particle size of AddaVax/OVA vaccine (*i*.*e*., 1, 2.5 and 5 mM). (F) The influence of antigen amount on the Z-average hydrodynamic particle size of AddaVax/OVA vaccine was investigated *with* AddaVax/OVA model vaccines with different amounts of OVA (*i*.*e*., 6, 12 or 50 *μ*g/100 *μ*L). Trehalose was employed as a stabilizer in all vaccine formulations at a concentration of 125 mg/mL. Vaccine formulations were frozen into thin films at a drum temperature of -100°C except for (D). Various antigens were loaded in the vaccine formulations at an antigen amount of 6 *μ*g/100 *μ*L except for (C) and (F). Formulations were prepared in citrate buffer (2.5 mM, pH 6.5) except for (E). \*\**p< 0*.*01, ns*: non-significant (*p>0*.*05*).

### 3.3. In vitro characterization of dry powders of AddaVax-adjuvanted influenza rHA vaccines

AddaVax/rHA vaccines (*i*.*e*., AddaVax/Y-2, AddaVax/J-4, AddaVax/Y-2/J-4, and AddaVax/Y-2/J-4/TJ-5) were prepared by dissolving one, two or three rHA proteins at a total rHA protein amount of 6 *μ*g in 50 *μ*L of citrate buffer (2.5 mM, pH 6.5) containing trehalose at a concentration of 250 mg/mL, which was then admixed with AddaVax (50 *μ*L). The vaccines were thin-film frozen at -100°C followed by sublimation to remove water. As shown in **Figures 5A and 6A**, the particle size and zeta potential values of all AddaVax/rHA vaccines did not significantly change (*p>0*.*05)* after subjected to TFFD and subsequent reconstitution. Additionally, subjecting the AddaVax/rHA vaccines to TFFD did not cause apparent aggregation nor degradation of rHA proteins based on SDS-PAGE data **(Figure 6B)**. Importantly, the hemagglutination activity of the rHA proteins were maintained after the AddaVax/rHA vaccines were subjected to TFFD **(Figure 6C)**. Overall, it appeared that subjecting protein antigens adjuvanted with AddaVax to TFFD did not comprise the integrity and activity of the antigens.

**Figure 6.**
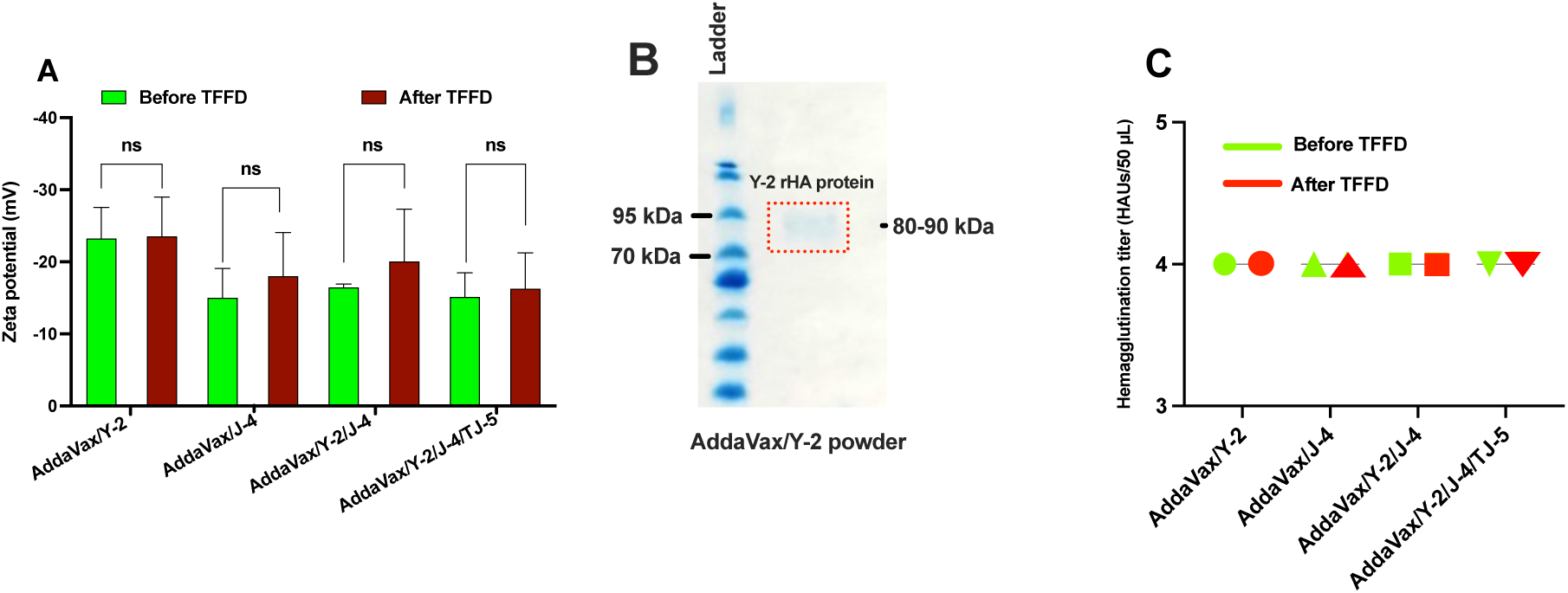
In vitro characterization of AddaVax-adjuvanted influenza rHA vaccines. (A) Zeta potential values of vaccine candidates determined by DLS. (B) SDS-PAGE analysis of Y-2 rHA proteins reconstituted from the vaccine dry powders. (C) Hemagglutination titers of various AddaVax/rHA vaccines before and after they were subjected to TFFD and subsequent reconstitution. Hemagglutination titer assay was repeated twice with the same results. *ns*: non-significant (*p>0*.*05*).

### 3.4. In vivo evaluation of AddaVax-adjuvanted rHA vaccines after subjected to TFFD

BALB/c mice were vaccinated twice at 4-week intervals to evaluate the immunogenicity of the dry powder AddaVax/rHA influenza vaccines, in comparison to the same vaccines that were not subjected to TFFD. Blood was collected 4 weeks after second vaccination and sera were analyzed for functional, neutralizing antibody titers against a panel of H1 and H3 viruses. Although mice are widely used for preclinical trials of influenza vaccines, many strains, including BALB/c, do not show disease symptoms after infection with clinical isolates of H3N2 [52]. For future studies, DBA/2J mice that have shown to be susceptible to infection by human clinical isolates of influenza will be used [53]. However, the animal studies demonstrated that the immunogenicity of AddaVax/rHA influenza vaccine dry powders was maintained and comparable to their liquid counterparts (**Figures 7-8**). Sera from mice that were not immunized (*i*.*e*., Mock group) (**Figure 7A**) or immunized with AddaVax alone (*i*.*e*., liquid AddaVax group) (**Figure 7B**) did not show any hemagglutination inhibition activity. Mice that were vaccinated with liquid or reconstituted dry powder of AddaVax/Y-2 vaccine candidate (Y-2 is an H1 rHA protein) produced HAI titers against currently circulating H1N1 influenza viruses, but not to pre-pandemic strains (**Figure 7C-D)**. A titer of 1:40 is accepted as a “protective” correlate of protection [54]. The HAI titers elicited by the liquid or reconstituted dry powder of AddaVax/Y-2 were not significantly different. Similarly, mice that were immunized with reconstituted dry powder of AddaVax/J-4 (J-4 is an H3 rHA protein) produced HAI titers against a broad range of H3N2 influenza viruses (**Figure 7E)**. Mice that were immunized with reconstituted dry powder of AddaVax-adjuvanted influenza vaccine candidates containing both H1 (*i*.*e*., Y-2) and H3 (*i*.*e*., TJ-5 and/or J-4) rHA proteins produced HAI titers against a broad range of H3N2 influenza viruses as well as currently circulating H1N1 influenza viruses (**Figure F-G**). There was also no significant difference between the HAI titers of reconstituted and liquid AddaVax/Y-2/J-4/TJ-5 vaccines **(Figure 7G *vs* 7H**), demonstrating that subjecting the AddaVax/rHA vaccines to TFFD did not affect the HAI activity of the antisera induced by the vaccines.

**Figure 7.**
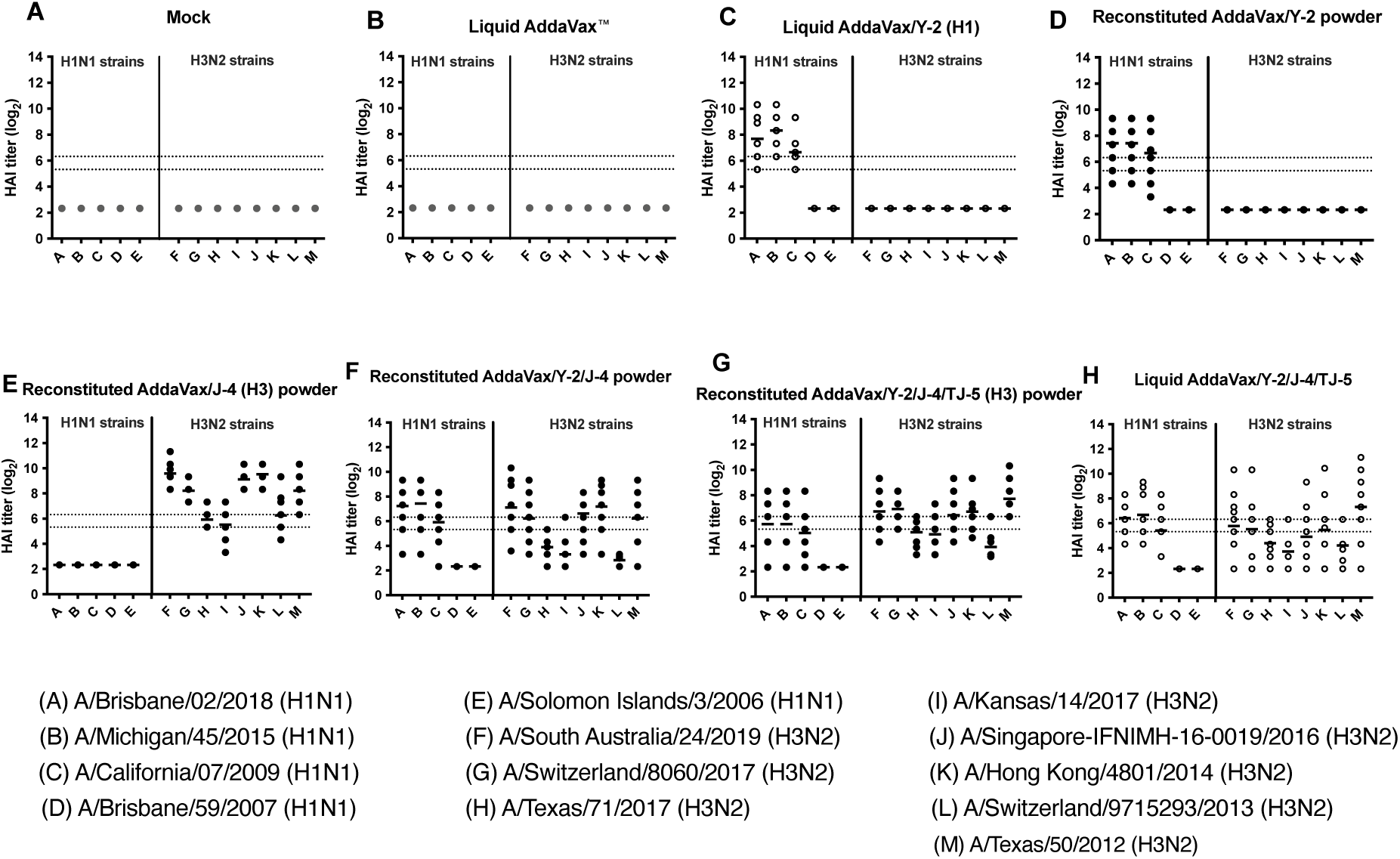
HAI serum antibody titers induced by TFFD formulated vaccines in mice. BALB/c mice (n=10) were vaccinated twice at 4-week intervals with AddaVax/rHA influenza vaccine candidates formulated as powders by TFFD. Control groups were vaccinated with the liquid formulation counterpart, AddaVax only, or mock. HAI titers were determined for individual mice, with mean values indicated, 4 weeks after boost. The x-axis represents the viruses used, with H1N1 strains on the left of the solid vertical line and H3N2 strains to the right. The bottom and top dashed horizontal lines indicate 1:40 and 1:80 titer, respectively.

To determine if the antibodies can block live virus infection, pooled sera from immunized mice were incubated with H1N1 influenza viruses (**Figure 8A-B**) or H3N2 viruses (**Figure 8C-E**). Then, the viruses’ ability to infect MDCK-SIAT1 cells was evaluted. Reconstituted dry powder AddaVax/Y-2 vaccine candidate and its liquid counterpart elicited similar neutralizing antibody titers against A/California/07/2009, though liquid AddaVax/Y-2 vaccine induced higher neutralizing antibody titers against A/Brisbane/2/2018. Reconstituted dry powder AddaVax/Y-2/J-4/TJ-5 vaccine and its liquid counterpart elicited similar neutralizing titers against A/Brisbane/2/2018, A/California/07/2009, A/Singapore-INFIMH-16-0019/2016, and A/Hong Kong/4081/2014, and the reconstituted dry powder vaccine elicited higher antibody titers to the A/Kansas/14/2017 than its liquid counterpart. Finally, 4 weeks after the second immunization, mice in all groups were intranasally challenged with A/Kansas/14/2017 (H3N2), but there was not any significant difference among any of the immunized groups in terms of weight loss (**Figure 3S**).

**Figure 8.**
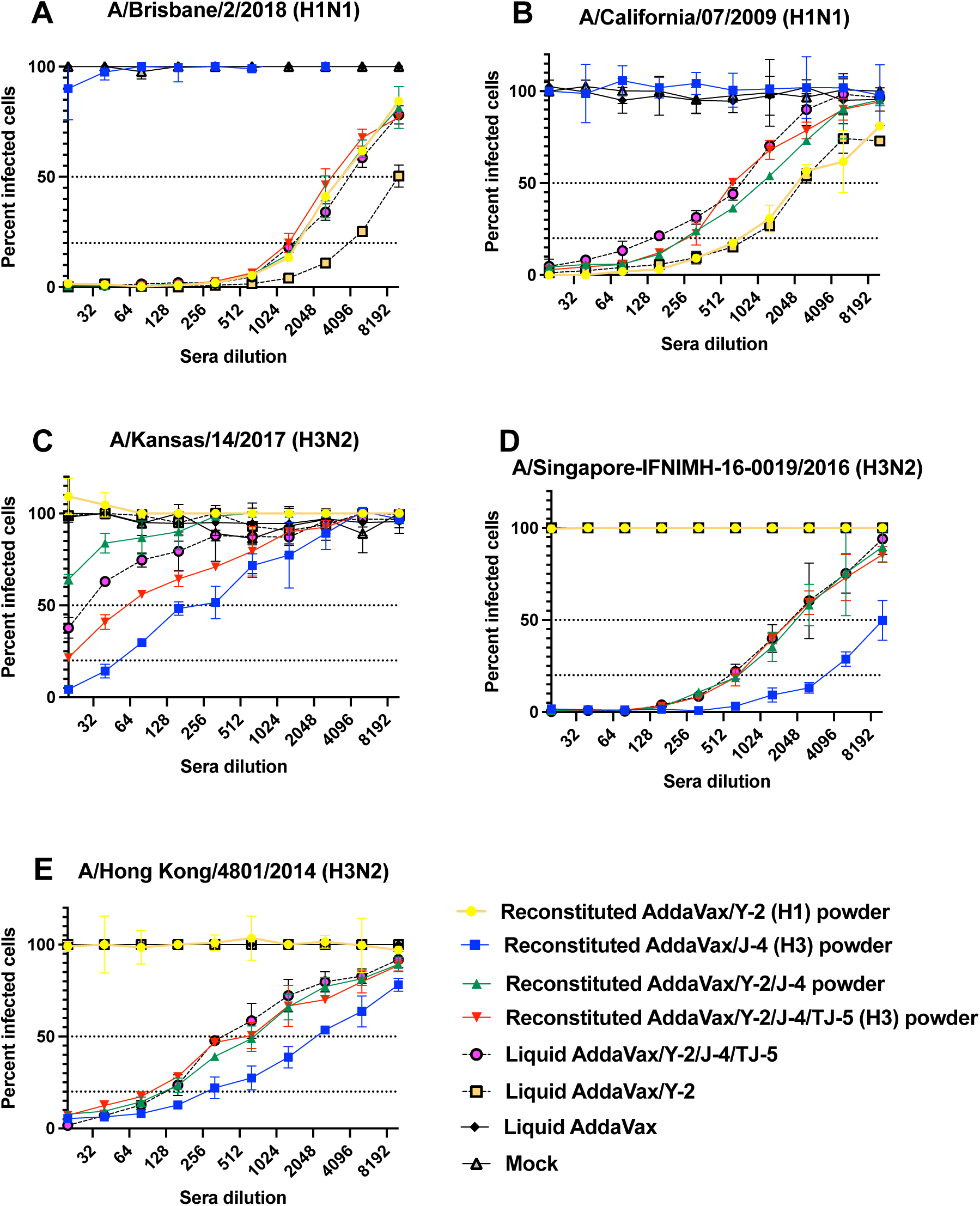
Neutralizing antibody titers induced by dry powders of AddaVax/rHA influenza vaccine candidates prepared using TFFD. FRA was done using pooled mouse sera 4 weeks after boost. BALB/c mice were vaccinated twice at 4-week intervals with reconstituted dry powders of AddaVax/rHA, liquid AddaVax/rHA vaccine formulations, reconstituted dry powder of AddaVax prepared by TFFD, or mock. Each graph represents a strain used for FRA against H1N1 (A-B) and H3N2 (C-E) viruses, with vaccine groups indicated on the right. The dotted dash lines represent 50% and 80% inhibition by sera compared to virus-only controls.

Vaccine particle size has been reported to affect the immunogenicity of the vaccine [18]. Data in Figures 7 and 8 showed that the slight mean particle size increase of the AddaVax-adjuvanted influenza vaccines after subjected to TFFD, though not significant, did not adversely affect their immunogenicity in mice, indicating that TFFD is a promising technology for the conversion of MF59-like nanoemulsion-adjuvanted vaccines into dry powders.

### 3.5. TFFD of Fluad Quadrivalent, an MF59-adjuvanted vaccine

Using various antigen and antigen combination, we have showed that TFFD can be applied to convert vaccines adjuvanted with AddaVax from liquid to dry powder with maintaining the immunogenicity of the vaccines. To confirm the applicability of TFFD to MF59-adjuvanted vaccines, commercially available Fluad Quadrivalent vaccine that contains MF59 was subjected to TFF at -100 °C and using trehalose at 125 mg/mL as a stabilizer. Upon sublimation and reconstitution, the particle size, PDI, and zeta potential of the vaccine as well as the integrity and function of the HA antigens in the vaccine were determined. As shown in **Figures 9A-B**, the particle size and PDI of the reconstituted vaccine increased slightly, but the zeta potential of the vaccine was maintained (**Figure 9C**). The hydrodynamic particle size increase can in part be due to the strong interaction of trehalose with the vaccine upon drying, and it is expected that further composition optimization can help minimize the size increase. Importantly, the integrity of the antigens (**Figure 9D**) and hemagglutination activity of the HA antigens in the vaccine (**Figure 9E**) remained unchanged, further confirming that TFFD can be applied to convert a vaccines adjuvanted with MF59 or AddaVax into dry powders. This is to our knowledge the first report of formulating an MF59-adjuvanted vaccine into a dry powder. As MF59 is mainly used in influenza vaccines, this report is an important step towards the development of a stable, universal influenza vaccine.

**Figure 9.**
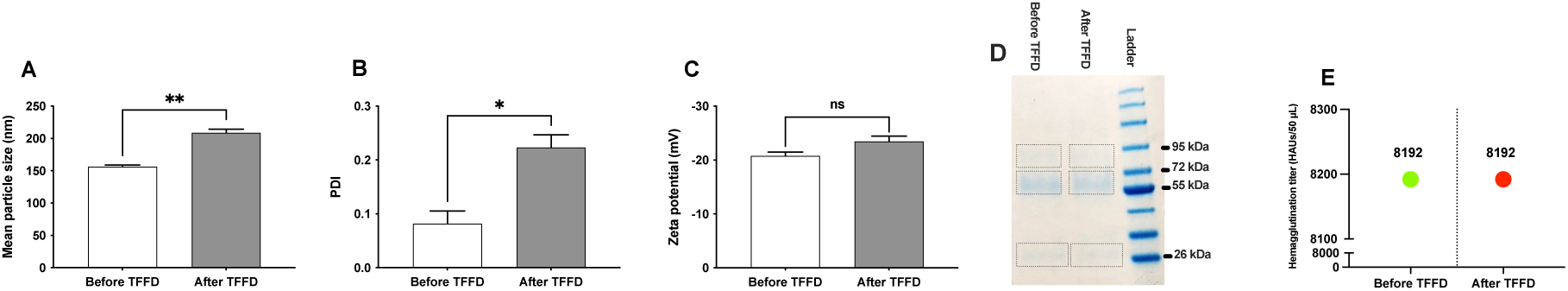
Characterization of Fluad Quadrivalent dry powder prepared by TFFD. (A) Mean particle size, (B) PDI and (C) zeta potential values of liquid vaccine *(i*.*e*., before TFFD) and reconstituted dry powder determined using DLS. (D) SDS-PAGE analysis. (E) Hemagglutination titers in HAUs/50 *μ*L. Hemagglutination was repeated twice (n = 2) with the same results were observed. Fluad Quadrivalent vaccine is adjuvanted with MF59 and contains the HA proteins of four influenza strains at 15 *μ*g/0.5 mL each. Trehalose dissolved in citrate buffer (2.5 mM, pH 6.5) was employed as a stabilizer at final concentration of 125 mg/mL. Trehalose (50 *μ*L) was mixed with 50 *μ*L of Fluad Quadrivalent and then the liquid vaccine formulation was frozen to thin films at -100°C. The vaccine dry powder was reconstituted in 100 *μ*L milli-Q water before characterization by DLS. Samples for SDS-PAGE analysis and hemagglutination assay were reconstituted in 50 *μ*L milli-Q water so that the HA content is 6 *μ*g/50 *μ*L (*i*.*e*., 60 g/0.5 mL) to facilitate the analysis. *\*p<0*.*05, **p<0*.*01, ns: non-significant (p<0*.*05)*.

### Conclusion

MF59 is a nanoemulsion adjuvant in FDA-approved human influenza vaccines. MF59-containing vaccines need cold chain for storage and transport, which may be avoided by converting the vaccines to dry powders. We report that TFFD can be applied to convert AddaVax, a preclinical grade equivalent of MF59, and vaccines containing MF59 or AddaVax from liquid to dry powders. The extent to which the particle size can be maintained was dependent on the antigen and the stabilizing excipient(s) used. Importantly, using monovalent, bivalent, and trivalent rHA antigens against H1 and/or H3 influenza viruses, we showed that subjecting the rHA vaccines adjuvanted with AddaVax to TFFD did not significantly affect the immunogenicity of the vaccines in a mouse model, pointing to the potential of developing a universal dry powder flu vaccine.

## Acknowledgments

This work was in part supported by Sponsored Research Agreements and Technology Validation Agreements from TFF Pharmaceuticals Inc. (to ROW, ZC, and TMR). KA is supported in part by a fellowship (GM 1105) from the Egyptian Ministry of Higher Education. TMR is supported, in part, by the University of Georgia and by the National Institute of Allergy and Infectious Diseases, a component of the NIH, Department of Health and Human Services, under contract 75N93019C00052. In addition, TMR is supported by the Georgia Research Alliance as an Eminent Scholar.

## Conflicts of interest

ZC, ROW, TMR report financial support by TFF Pharmaceuticals, Inc. ZC reports a relationship with TFF Pharmaceuticals, Inc. that includes: equity or stocks and funding grants. ROW reports a relationship with TFF Pharmaceuticals, Inc. that includes: consulting or advisory, equity or stocks, and funding grants. HX, CM, and SS report a relationship with TFF Pharmaceuticals, Inc. that includes: consulting or advisory. ZC and ROW have a patent “Dry solid aluminum adjuvant-containing vaccines and related methods thereof” pending to TFF Pharmaceuticals, Inc. ZC, ROW, KA, HX, and CM have a patent “Dry powder compositions of oil-in-water (O/W) emulsion adjuvanted vaccines” pending to UT Austin. DJC is a paid consultant for TFF Pharmaceuticals, Inc. TMR is a member of the TFF Pharmaceuticals, Inc. Scientific Advisory Board.

