## Supplemental tables and figures for "Formulation of Dry Powders of Vaccines Containing MF59 or AddaVax by Thin-Film Freeze-Drying"

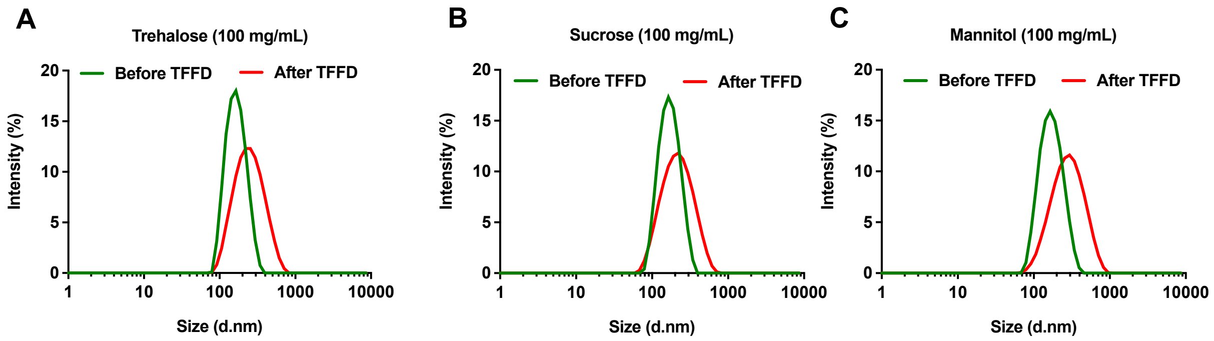


**Figure 1S.** Particle size distribution of AddaVax/OVA incorporating different stabilizers at 100 mg/mL. Stabilizers were dissolved in citrate buffer (2.5 mM, pH 6.5) before mixing with AddaVax™ (50 µL) and OVA antigen (6 µg). Liquid AddaVax/OVA vaccine formulations were then frozen into thin films at drum temperature of -100ºC followed by water sublimation.

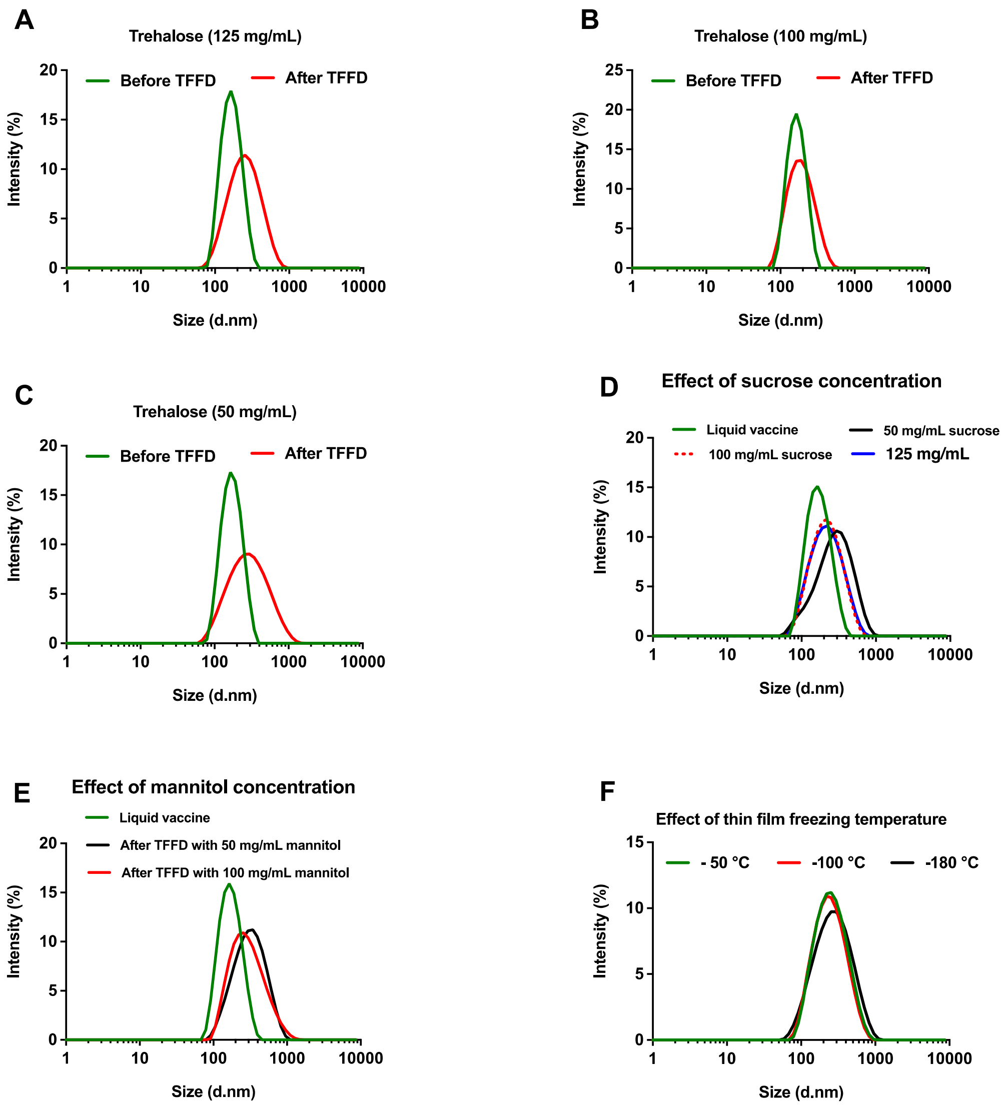


**Figure 2S. Effect of stabilizing excipient concentration and TFF temperature on the Z-average particle size of TFFD-processed powders of aAddaVax/OVA model vaccine.** Trehalose was employed as a stabilizing agent at different concentrations including (A) 125 mg/mL, (B) 100 mg/mL, and (C) 50 mg/mL. (D) Sucrose was also screened at 50, 100 and 125 mg/mL and (E) mannitol was screened at 50 and 100 mg/mL. (F) Effect of TFF temperature on particle size distribution of the model vaccine.

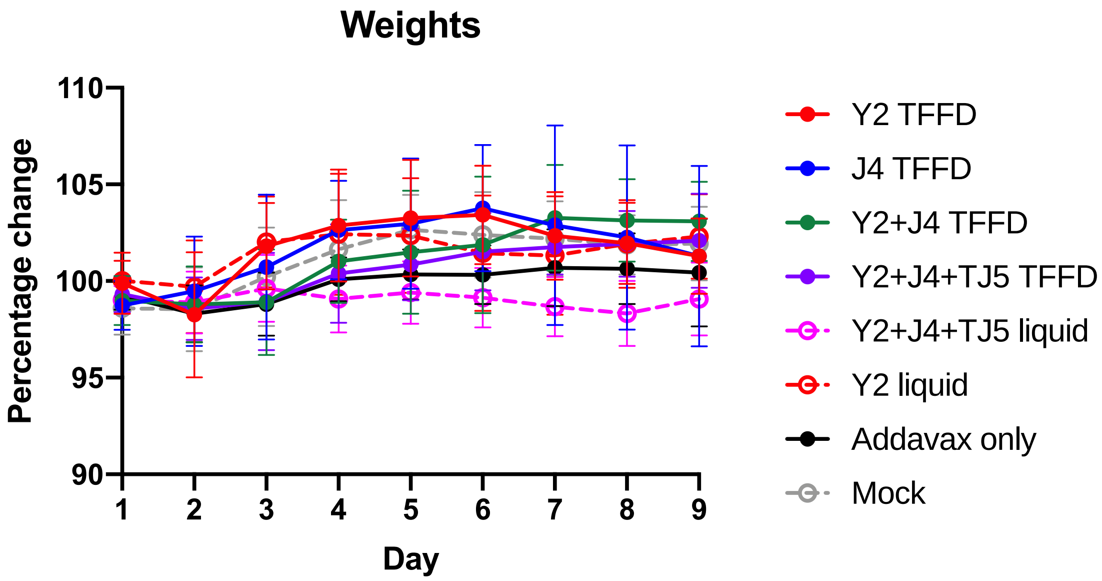


**Figure 3S.** Mouse weights after H3N2 challenge.
